# Anti- *Leishmania*-activated C-kinase monoclonal antibody immunohistochemical technique for cutaneous leishmaniasis diagnosis

**DOI:** 10.64898/2025.12.14.694232

**Authors:** Amanda Desirée Lopes Santos, Marcelo Antônio Pascoal-Xavier, Vanessa Peruhype-Magalhães, Marcela Tavares Caldas Eller, Francislane Ramalho Varella Pereira, Maryana Prates Rodrigues, Maria Nilza Pereira de Brito, Bruno Oliveira Souza e Silva, Thadeu Ramalho da Silva, Daniel Moreira de Avelar, Mariana Lourenço Freire, Edward Oliveira

## Abstract

This study aimed to develop and validate an immunohistochemistry (IHC) technique that employs an anti- *Leishmania-*activated C kinase (LACK) antigen (anti-LACK) monoclonal antibody (mAb) for the diagnosis of cutaneous leishmaniasis (CL). The *Leishmania braziliensis* LACK sequence (A4HGX7_LEIBR) was analyzed for B cell epitope prediction and three antigenic regions were selected to design a multiepitope antigen. The corresponding gene was back translated, synthesized and cloned into the pET28a (+) plasmid. Recombinant *Leishmania*-LACK was expressed and used as an immunogen for mAb production via somatic hybridization. The ability of the anti-LACK mAb to recognize native LACK and *Leishmania (Leishmania) amazonensis*, *Leishmania (Viannia) braziliensis*, and *Leishmania (Viannia) guyanensis* amastigotes was confirmed in total extracts and skin histological sections from experimentally infected hamsters using Western blotting and IHC techniques, respectively. The anti-LACK IHC was then validated on skin lesion samples from 104 suspected CL patients attending the outpatient clinic of the Municipal Polyclinic of Teófilo Otoni between 2019 and 2020, using kDNA-PCR as reference test. The diagnostic performance of anti-LACK IHC was compared to direct (DE) and histopathological (HE) examinations. The anti-LACK mAb successfully recognized native LACK in soluble antigens and detected amastigotes of the three *Leishmania* species in histological sections from experimentally infected hamsters. Moreover, anti-LACK IHC presented sensitivity of 59.3% (95% CI: 40 – 83.7), specificity of 98% (95% CI: 89.5– 99.7), and accuracy of 77.9% (95% CI: 61.9 – 96.8), higher than that presented by DE and HE techniques, although differences were not statistically significant. Agreement analysis yielded a Kappa index of 0.56 (95% CI: 0.42-0.71) considering CL cases compared with kDNA-PCR results. The combination of DE or HE with anti-LACK IHC increased diagnostic accuracy to 83.7% (*p*: 0.48) and 79.8% (*p*: 0.33), respectively. In this initial study, the anti-LACK IHC detected the main *Leishmania* species causing CL in Brazil, yet further improvement promises enhanced diagnostic performance of this technique.

## INTRODUCTION

Cutaneous leishmaniasis (CL) remains endemic in several regions worldwide, including the Americas, Africa, the Eastern Mediterranean, and Southeast Asia, with approximately 272,098 new cases reported in 2023 [1]. Although the Americas experienced an 8% decrease in the number of cases, the Pan American Health Organization (PAHO) still reported 34,954 cases in 2023 [2], of which 12,910 occurred in Brazil [2]. The main causal species in Brazil are *Leishmania* (*Viannia*) *braziliensis*, *Leishmania* (*Leishmania*) *amazonensis* and *Leishmania* (*V.*) *guyanensis* [3]. In addition, other species such as *Leishmania* (*V*.) *naiffi*, *Leishmania* (*V*.) *shawi*, *L*. (*V*.) *lindenberg*, *Leishmania* (*V*.) *lainsoni* and *L*. (*L*) *infantum* have also been identified, mainly in the North, Northeast and VL-endemic regions of Brazil [4–8]. The high incidence and wide geographical distribution of CL, together with the risk of progression to chronic forms, make it a significant public health challenge.

Accurate laboratory diagnosis is essential for the clinical management of CL, particularly considering the toxicity of available treatments and the clinical similarities between CL and other diseases such as sporotrichosis, vascular ulcers, leprosy, paracoccidioidomycosis, skin cancer, and deep mycoses [9]. In clinical settings, diagnosis is primarily based on direct examination (DE), culture, and, when available, polymerase chain reaction (PCR). However, these techniques often show limitations related to low sensitivity or technical complexity, which restricts their use in many endemic regions. Improving the accessibility and performance of diagnostic tools is a priority outlined in the United Nations 2030 Agenda, which aims to eliminate neglected tropical diseases by the end of the decade [10].

In this context, immunohistochemistry (IHC) has emerged as a promising diagnostic alternative as it allows for the detection of parasite antigens in tissue samples through the use of specific antibodies [11]. Despite significant advancements in IHC methodologies, its application to CL diagnosis remains limited. Most studies have used hyperimmune sera or biotin-based detection systems [12–17]. Only a few studies have reported the use of monoclonal antibodies (mAbs) in IHC for CL diagnosis. Shirian et al. (2014) achieved over 94% sensitivity for *Old World* species [18], while Freire et al. (2022) reported up to 85.7% sensitivity for *New World* species [19].

The development of mAbs targeting highly expressed parasite antigens may enable IHC-based detection of *Leishmania* spp., even in samples with low parasite loads. Given this background, the present study aimed to develop and validate an IHC technique employing a mAb against the *Leishmania*-activated C kinase (LACK) antigen, combined with a polymer-based detection system, for the diagnosis of CL.

## MATERIALS AND METHODS

### Antigenic Target

LACK is a highly conserved protein among *Leishmania* species, expressed in both amastigote and promastigote stages. It plays essential roles in temperature tolerance, parasite survival, and virulence [20,21], and is also involved in the early stages of the host immune response [22]. LACK is predominantly localized in the cytoplasm near the kinetoplast but has also been detected on the parasite surface, possibly due to the reassociation of its secreted form [23–25].

Given its multiple biological functions, particularly in temperature adaptation, translation, and virulence [20,21], LACK may act as a molecular coordinator linking environmental cues to amastigote development and/or replication. Based on these observations, it is plausible that threshold levels of LACK regulate the expression of one or more virulence-associated proteins during mammalian infection, enhancing parasite fitness during host cell invasion, differentiation, and replication under hostile intracellular conditions.

Due to its role in *Leishmania* immunopathogenesis, LACK has been extensively studied as a vaccine target. Kelly et al. [20] demonstrated that deletion of the *LACK* gene in *L. major* significantly reduced parasite burden in vivo, reinforcing its contribution to virulence. Furthermore, an IHC assay employing an anti-LACK mAb achieved 100% sensitivity for detecting amastigotes in visceral leishmaniasis tissue samples (bone marrow and liver), underscoring the antigen’s diagnostic potential. However, the performance of anti-LACK IHC has not yet been evaluated for the diagnosis of CL.

### Production of the Recombinant LACK Antigen

The amino acid sequences of the LACK protein from *L. braziliensis* (ID: A4HGX7_LEIBR), *L. amazonensis* (ID: Q95NJ3), and *L. guyanensis* (ID: A0A1E1J0L7) were compared using the Clustal Omega program. The *L. amazonensis* sequence showed ten amino acid differences at positions 43, 58, 99, 148, 183, 231, 237, 254, 256, and 277, resulting in 96.79% overall identity.

Subsequently, the *L. braziliensis* LACK sequence was analyzed using BCPRED and Bepipred 2.0 for B-cell epitope prediction. Three antigenic regions were selected (Fig S1), reverse-translated, and used to design a synthetic gene containing a start codon, *BamHI* and *HindIII* restriction sites, and a stop codon. Sequence design and optimization were performed using BioEdit v7.2.5 (Raleigh, NC, USA). The final 645-bp construct, including a 630-bp open reading frame encoding a 210-amino-acid protein (23.1 kDa), was cloned into the pET28a(+) vector for heterologous expression in *Escherichia coli* BL21 Star cells.

Following heat-shock transformation, recombinant clones were screened by *BamHI/HindIII* digestion and cultured in Luria–Bertani (LB) medium supplemented with kanamycin (30 µg/mL). Protein expression was performed as described by Freire et al. (2022) [19]: an 80-mL pre-culture was expanded to 2 L of LB medium, induced with 0.75 mM IPTG at OD₅₉₀ 0.6–0.8, and incubated overnight at 16 °C with shaking (200 rpm). Bacterial cells were harvested (10,000 × g, 20 min) and lysed under denaturing conditions to isolate inclusion bodies.

Cells were resuspended in lysis buffer (50 mM Tris-HCl, 500 mM NaCl, 0.2 mM EDTA, 3% sucrose, 1% Triton X-100, 200 µg/mL lysozyme, 1 mM PMSF, 20 µg/mL DNase I; pH 8.0) and sonicated on ice for five 30-s cycles (VC-750, Sonics Vibra-Cell, Sonics & Materials, Newton, CT, USA). Inclusion bodies were pelleted (13,000 × g, 40 min), washed with Tris–urea buffer (50 mM Tris-HCl, 3 M urea, 0.2 mM EDTA, 500 mM NaCl, pH 8.0), re-sonicated, and centrifuged again. Solubilization was achieved in phosphate buffer (10 mM) containing NaCl (200 mM), Tris-HCl (10 mM), guanidine-HCl (6 M), and β-mercaptoethanol (10 mM).

Recombinant LACK was purified using Ni Sepharose HP resin in Poly-Prep® chromatography columns (Bio-Rad Laboratories Inc. Hercules, CA, USA). Protein expression and purity were confirmed by SDS-PAGE. For Western blotting, proteins were transferred to nitrocellulose membranes and probed with anti-6×His antibody (1:3,000; Thermo Fisher) followed by enhanced chemiluminescence (ECL) detection (Cytiva, Buckinghamshire, ENG). Images were acquired using an ImageQuant LAS 4000 system (Cytiva, Upsala, SWE).

### Production of the Monoclonal Antibody

Two five-week-old female BALB/c mice were subcutaneously immunized with 20 µg of recombinant LACK (rLACK) protein emulsified in Freund’s complete adjuvant, followed by four booster doses (20 µg each) in Freund’s incomplete adjuvant at 15-day intervals. Control mice received saline with adjuvant following the same immunization schedule. Blood samples were collected from the submandibular vein prior to each immunization.

Serum IgG antibody levels were determined by indirect ELISA using rLACK as the coating antigen. Microplates (Nunc Maxisorp™) were coated with 1 µg/mL rLACK in carbonate–bicarbonate buffer (pH 9.6) and incubated for 1 h at 37 °C, followed by overnight incubation at 2–8 °C. After washing with PBS containing 0.05% Tween-20 (PBS-T20), wells were blocked with 5% skim milk in PBS-T20 for 2 h at 37 °C. Serum samples (1:100 dilution in 1% PBS-T20-milk) were added and incubated for 1 h at 37 °C. Plates were washed and incubated with peroxidase-conjugated anti-mouse IgG (1:20,000; Thermo Fisher Scientific), followed by addition of TMB substrate. After 5 min, the reaction was stopped with 1 N H₂SO₄, and absorbance was measured at 450/620 nm using a Varioskan LUX reader (Thermo Fisher Scientific).

Following confirmation of specific IgG production, mice received an intraperitoneal booster dose of rLACK. Three days later, the animals were euthanized by ketamine (500 mg/kg) and xylazine (50 mg/kg) anesthesia, and spleens were collected for somatic cell fusion with *Sp2/0-IL6* myeloma cells (3.5 × 10⁷ splenocytes: 7 × 10⁶ myeloma cells), as described by Freire et al. (2022)[19]. Fusion was performed using PEG-DMSO (Hybri-Max), and cells were washed and cultured in DMEM supplemented with 20% fetal bovine serum (FBS) and 1% Penicillin–Streptomycin for 24 h at 37 °C under 5% CO₂. Subsequently, selective HAT medium (2%) was added to 96-well plates previously seeded with murine peritoneal macrophages.

After 10–14 days, hybridoma supernatants were screened by ELISA. The highest-producing clone (absorbance ≥ 1.20) was subcloned by limiting dilution to obtain a monoclonal population. The selected clone was expanded, and culture supernatants containing the anti-LACK mAb were collected weekly. Supernatants were concentrated using the Amicon® Stirred Cell system with Diaflo ultrafilters and purified sequentially on HiTrap Blue HP and rProtein A FF affinity columns (Cytiva).

Purity of the anti-LACK mAb was assessed by 15% SDS-PAGE. Antigen recognition was confirmed by Western blot analysis using 1 µg of rLACK as antigen and anti-LACK mAb at a 1:100 dilution.

### Recognition of Native LACK Antigen in Soluble Leishmania spp. Antigens

The ability of the anti-LACK mAb to recognize the native antigen in total protein extracts from the main *Leishmania* species causing CL in Brazil was evaluated by Western blotting. Total extracts of *L. (L.) amazonensis*, *L. (V.) braziliensis*, and *L. (V.) guyanensis* (as described by Freire et al., 2022 [19]) were resolved by 15% SDS-PAGE.

Separated proteins were electrotransferred onto nitrocellulose membranes (Amersham Protran, 0.45 µm; Cytiva) [26]. Membranes were blocked in 5% skim milk in PBS–Tween 20 (0.05%) and incubated with the anti-LACK mAb (1:3,000) for 1 h at room temperature. After washing, membranes were incubated with horseradish peroxidase (HRP)–conjugated anti-mouse IgG (Sigma-Aldrich, St. Louis, MO, USA) for 1 h. Immunoreactive bands were detected using ECL™ Prime Western Blotting Detection Reagent (Cytiva), and images were acquired using the ImageQuant LAS 4000 system (Cytiva).

### Detection of Amastigote Forms of Leishmania spp. in Histological Sections from Experimentally Infected Animals

The ability of the anti-LACK mAb to detect *Leishmania* amastigotes in tissue was evaluated by performing IHC on histological sections of dermal lesions from hamsters (*Mesocricetus auratus*) experimentally infected with the three main *Leishmania* species responsible for CL in Brazil.

Five-week-old male golden hamsters (*Mesocricetus auratus*) were infected with *L. (L.) amazonensis* (IFLA/BR/67/PH8), *L. (V.) braziliensis* (MHOM/BR/75/M2903), or *L. (V.) guyanensis* (MHOM/BR/75/M4147). Each animal received a subcutaneous inoculation of 1 × 10⁶ promastigotes/mL into the dorsal surface of the right hind paw. Skin lesion fragments were collected 30–40 days post-infection and fixed in 10% neutral buffered formalin (pH 7.2).

Tissue sections of 4-µm thickness were prepared using a rotary microtome, mounted on positively charged glass slides, and subjected to IHC using the anti-LACK mAb. A biotin-free, polymer-based detection system conjugated to alkaline phosphatase (Bond Polymer Refine Red Detection, Leica Microsystems, Newcastle, UK) was employed, resulting in red/burgundy staining of parasite amastigotes.

### Standardization of the IHC Protocol Using Anti-LACK Monoclonal Antibody

IHC using the anti-LACK mAb (anti-LACK IHC) was standardized with five paraffin-embedded skin biopsy samples from patients with confirmed CL, previously diagnosed by microscopy and quantitative PCR (qPCR) targeting kinetoplast DNA (kDNA). The procedure was adapted from Freire et al. (2022)[19], with minor modifications.

Briefly, 4-µm tissue sections were mounted on ImmunoSlides (EasyPath Diagnósticos, Indaiatuba, São Paulo, BRA) and incubated overnight at 56 °C. Dewaxing, rehydration, and antigen retrieval were performed using Trilogy® (Cell Marque, Rocklin, CA, USA) diluted 1:100 and heated in a steamer for 30 min at approximately 90 °C. The IHC protocol consisted of the following steps: (I) Blocking with 5% skim milk in PBS for 30 min; (II) Incubation with anti-LACK mAb for 60 min; (III) Application of post-primary alkaline phosphatase (AP)–conjugated antibody for 30 min; (IV) Incubation with polymer-based AP reagent for 30 min; (V) Chromogen development using the Red reagent for 3 min; and (VI) Counterstaining with hematoxylin for 3 min.

Blocking time, antibody dilution, and chromogen incubation parameters were optimized to achieve high specificity with minimal background staining (Supplementary Table S1).

### Validation of the Immunohistochemistry Technique with Human Samples

#### Participants and Samples

Patients with clinical suspicion of CL who attended the Municipal Polyclinic in Teófilo Otoni, Brazil, between 2019 and 2020 were included in this study. For each patient, the lesioned area was cleaned with chlorhexidine and anesthetized with 2% lidocaine. Two 4-mm punch biopsies were then obtained from the active border of each ulcerative lesion.

One biopsy specimen was used to prepare imprints on glass slides for direct examination (DE), after which it was stored in microtubes at −20 °C for DNA extraction and PCR analysis. The second biopsy specimen was fixed in 10% neutral buffered formalin and processed for histopathological (HE) and immunohistochemical (anti-LACK IHC) examinations. All samples were anonymized, and laboratory personnel were blinded to the clinical status of the samples.

#### Direct Examination (DE)

Direct microscopy was performed at the *Laboratório Macroregional de Teófilo Otoni*, part of the *Rede Estadual de Laboratórios de Saúde Pública do Estado de Minas Gerais*, as part of routine CL diagnosis. Each biopsy specimen was gently pressed eight times against a glass slide to obtain tissue imprints, which were air-dried, fixed in methanol, and stained with Giemsa. Slides were examined under a light microscope at 100× magnification. Samples were considered positive when amastigote forms were visualized in the imprints.

#### Histopathological Examination (HE)

Skin lesion fragments were fixed in 10% buffered formalin (pH 7.2) for at least 24 h. The tissues were then dehydrated, cleared, and embedded in paraffin using an automatic tissue processor (PT05 TS, Lupetec, São Carlos, SP, BRA) and an inclusion center (CI 2014, Lupetec). Sections of 4 µm were cut, mounted on glass slides, and stained with hematoxylin-eosin. Slides were examined under 10×, 40×, and 100× magnifications using a Zeiss microscope (Hallbergmoos, GER) for the presence of *Leishmania* amastigotes [27]. Samples were considered positive when amastigote forms were visualized in the sections.

#### Polymerase Chain Reaction (kDNA PCR)

Total DNA was extracted from patient biopsies using the DNeasy Blood and Tissue Kits (Qiagen, Hilden, Germany), according to the manufacturer’s instructions. Genomic DNA from *L. braziliensis* promastigotes cultured in NNN/LIT biphasic medium supplemented with 20% inactivated fetal bovine serum (FBS) and 1% Penicillin–Streptomycin was used as a positive control. DNA concentration and purity were measured with a NanoDrop spectrophotometer (Cytiva, Chicago, IL, USA).

PCR amplification targeted a 120-bp fragment of the *Leishmania* kinetoplast minicircle DNA using primers 150 (sense) 5′-(C/G)(C/G)(G/C)CC(C/A)CTAT(T/A)TTACACCAACCCC-3′ and 152 (antisense) 5′-GGGGAGGGGCGTTCTGCGAA-3′ [28,29]. Each 20 µL reaction contained 2 µL 10× PCR buffer, 1.5 mM MgCl₂, 0.5 µM of each primer (Integrated DNA Technologies, Coralville, IA, USA), 200 µM dNTPs, 20 ng of template DNA, and ultrapure water.

Amplifications were carried out in a thermocycler (ProFlex PCR System, Thermo Scientific, Foster City, CA, USA) under the following conditions: initial denaturation at 94 °C for 4 min; 35 cycles of 94 °C for 30 s, 60 °C for 30 s, and 72 °C for 30 s; and a final extension at 72 °C for 10 min. PCR reactions containing no DNA template and *L. braziliensis* DNA were used as negative and positive controls, respectively. Amplicons were resolved by 1% agarose gel electrophoresis stained with ethidium bromide, and bands were visualized using the ImageQuant LAS 4000 system (Cytiva).

#### Anti-LACK IHC Technique

The optimized anti-LACK IHC protocol was performed using the Bond Polymer Refine Red Detection system (Leica Microsystems, Newcastle, ENG) for visualization of the antigen-antibody complex. Dewaxing, rehydration, and antigen retrieval were performed in a 1:100 dilution of Trilogy® solution (Cell Marque, Rocklin, CA, USA) for 30 min at 90 °C in a steamer. Non-specific binding was blocked with PBS containing 5% skim milk for 30 min at room temperature (RT).

Slides were incubated with the anti-LACK mAb (1:50) diluted in PBS containing 1% bovine serum albumin (BSA) and 0.1% sodium azide for 60 min at RT. After washing in Tris buffer (5 mM Tris base, 140 mM NaCl, pH 7.6), slides were incubated with the post-primary alkaline phosphatase (AP) antibody for 30 min at RT, followed by incubation with the polymer-based AP reagent for another 30 min.

The chromogenic reaction was developed using the Red reagent mixture (Part A 1:10, Part B 1:50, Part C 1:50 in Part D) for 3 min. Slides were rinsed in distilled water, counterstained with hematoxylin for 3 min, mounted with Entellan® (Merck KGaA, Darmstadt, GER), and examined under 10×, 40×, and 100× magnifications using a Zeiss microscope to detect *Leishmania* amastigotes.

#### Database and Statistical Analyses

Diagnostic performance was analyzed using MedCalc software version 15.0 (MedCalc Software, Ostend, BEL). Comparisons between diagnostic methods were performed using the chi-square test, with a significance level of 5%. Agreement between tests was assessed using Cohen’s Kappa coefficient, interpreted according to Landis and Koch (1977): <0 (poor), 0.00–0.20 (slight), 0.21–0.40 (fair), 0.41–0.60 (moderate), 0.61– 0.80 (substantial), and 0.81–1.00 (almost perfect).

## RESULTS

### Expression and Purification of Recombinant LACK (rLACK)

The 210–amino acid sequence derived from *L. braziliensis* LACK was back-translated and optimized for expression in a prokaryotic system. Successful transformation of *E. coli* BL21 Star cells with the synthetic *LACK* gene (630 bp) was confirmed by restriction digestion analysis (Fig 1a).

**Fig 1.**
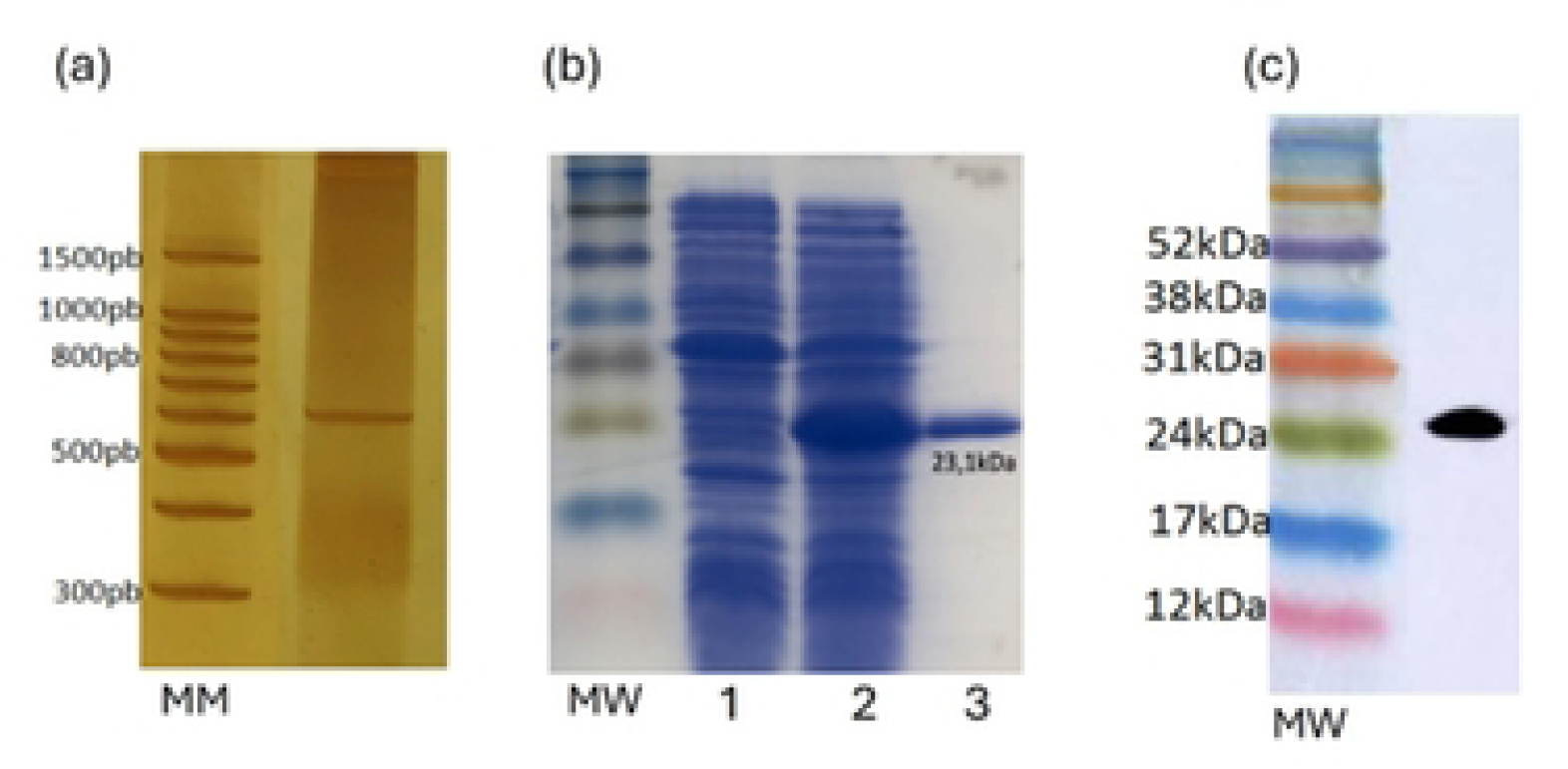
Production of recombinant LACK (rLACK) Production and characterization of rLACK: (a) Polyacrylamide gel showing silver-stained synthetic LACK gene fragment; (b) 15% SDS-PAGE of bacterial lysates before (1) and after (2) IPTG induction and purified 23.1 kDa rLACK (3); (c) Western blotting using anti-6x-His-tag mAb against rLACK. MM: 100 bp DNA Ladder (Promega, USA); MW: Amersham ECL Rainbow Markers (Cytiva, Chicago, IL, USA).

Protein expression and purification were verified by 15% SDS–PAGE, which demonstrated a distinct band corresponding to the expected molecular weight of the recombinant LACK (rLACK) protein after IPTG induction (Fig 1b). The identity of the expressed protein was further confirmed by Western blotting using an anti–6×His mAb, which detected a single band of approximately 23 kDa (Fig 1c).

### Production of the anti-LACK monoclonal antibody

A marked anti-LACK antibody response was observed in immunized mice prior to the third immunization, with antibody titers remaining high after the fifth dose, as determined by ELISA (Fig 2a). The specificity of the anti-LACK mAb for the recombinant antigen (rLACK) was confirmed by Western blotting (Fig 2b). Moreover, the antibody successfully recognized the native LACK antigen in soluble *Leishmania* extracts from *L. (L.) amazonensis* (SLaA), *L. (V.) braziliensis* (SLbA), and *L. (V.) guyanensis* (SLgA) (Fig 2c).

**Fig 2.**
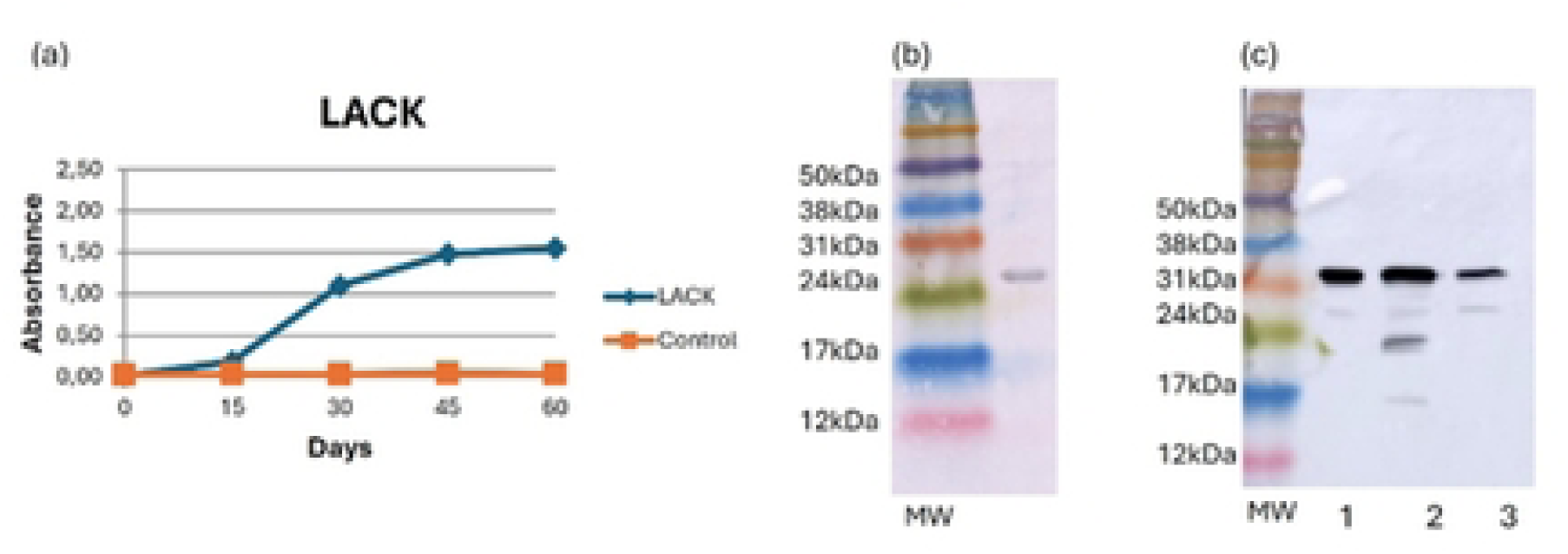
Production and characterization of the anti-LACK monoclonal antibody (mAb) Anti-LACK monoclonal antibody production and characterization procedures: (a) Kinetics of IgG antibody production in mice immunized with rLACK or receiving saline containing Freund’s adjuvant, measured by ELISA before each immunization. (b) Western blot showing recognition of rLACK by the specific anti-LACK mAb. (c) Western blot demonstrating reactivity of the anti-LACK mAb against soluble *Leishmania* antigens from *L. (L.) amazonensis* (1), *L. (V.) braziliensis* (2), and *L. (V.) guyanensis* (3). MW: Amersham ECL Rainbow molecular weight markers (Cytiva, Chicago, IL, USA).

### Recognition of Leishmania spp. in lesions from experimentally infected animals

IHC using the anti-LACK mAb successfully detected and labeled amastigote forms of *L. (L.) amazonensis*, *L. (V.) braziliensis*, and *L. (V.) guyanensis* in skin lesion samples from experimentally infected hamsters. Detection was performed using the Bond Polymer Refine Red system, resulting in clear visualization of the parasites within tissue sections (Fig 3).

**Fig 3.**
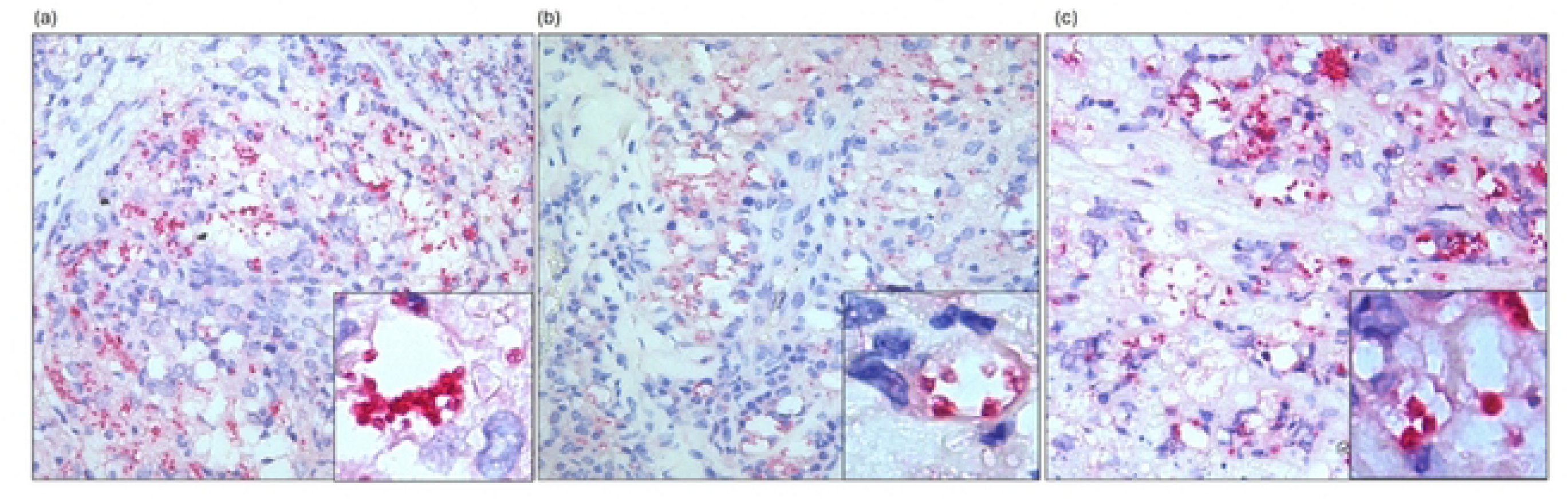
Immunohistochemical detection of *Leishmania* amastigotes in experimentally infected hamsters. Histological sections of skin lesions from golden hamsters infected with *L. (L.) amazonensis* (a), *L. (V.) braziliensis* (b), and *L. (V.) guyanensis* (c), showing positive immunolabeling of amastigote forms using the anti-LACK monoclonal antibody (mAb). The antigen-antibody reaction was visualized with the Bond Polymer Refine Red Detection system.

### Validation of anti-LACK IHC for cutaneous leishmaniasis diagnosis

A total of 104 participants who attended the Municipal Polyclinic in Teófilo Otoni, Brazil, between 2019 and 2020 were included in this study. The kDNA-PCR assay was used as the reference standard for classifying individuals as CL cases or non-CL cases. Among the participants, 54 were PCR-confirmed CL cases, while 51 were PCR-negative individuals with suspected alternative etiologies.

Of the confirmed CL cases, 11/54 participants resided in the urban area of Teófilo Otoni, while the remaining 43/54 lived in the surrounding regions, including rural communities and districts. This group comprised 23 females (median age: 42 years, range 23–49) and 31 males (median age: 53 years, range 41–61). Thirty-nine participants presented a single lesion (0.24–15.3 mm²), six had two lesions (0.09–3.3 mm²), four had three lesions (0.19–10.5 mm²), and three had four lesions (0.35–10.4 mm²). Two patients exhibited multiple lesions for which the total area was not measured. The duration of illness ranged from 0.67 to 12 months. All CL cases were treated with meglumine antimoniate (Glucantime®, Sanofi-Aventis Farmacêutica Ltda., Suzano, SP, BRA) in accordance with the Brazilian Ministry of Health guidelines (Brasil, 2017).

The diagnostic performance of the anti-LACK IHC was compared with HE and DE. The anti-LACK IHC assay showed the highest sensitivity (59.3%), followed by DE (53.7%) and HE (42.6%). Specificity exceeded 95% for all three methods. The anti-LACK IHC also achieved the highest overall accuracy (77.9%), followed by DE (74%) (Table 1). However, no statistically significant differences were observed among the performances of the three diagnostic tests.

**Table 1.** Diagnostic performance parameters of anti-LACK monoclonal antibody immunohistochemistry (anti-LACK IHC), direct examination (DE) and histopathological examination (HE)

| Diagnostic tests | Sensitivity (%) [CI95%]<br>n | Specificity (%) [CI95%]<br>n | Accuracy (%) [CI95%]<br>n |
| --- | --- | --- | --- |
| anti-LACK IHC | 59.3 [40.5-83.7]<br>(32/54) | 98 [89.5 – 99.7]<br>(49/50) | 77.9 [61.9-96.8]<br>(81/104) |
| DE | 53.7 [36 – 77.1]<br>(29/54) | 98 [89.5 - 99.6]<br>(49/50) | 75.0 [59.3- 93.6]<br>(78/104) |
| HE | 42.6 [27- 63.9]<br>(23/54) | 96 [86.5 – 99]<br>(48/50) | 68.3 [53.3-86.1]<br>(71/104) |

The agreement between PCR–kDNA, anti-LACK IHC, and other diagnostic techniques in classifying CL cases and non-cases is presented in Table 2. The Kappa coefficient values ranged from 0.38 to 0.56, indicating fair to moderate agreement. The lowest concordance was observed between PCR–kDNA and HE, whereas the highest agreement was found between PCR–kDNA and the anti-LACK IHC assay.

**Table 2.** Agreement analysis of the results of the diagnostic techniques applied to CL cases and non-CL cases.

|  | anti-LACK IHC |  |  | DE |  |  | HE |  |  |
| --- | --- | --- | --- | --- | --- | --- | --- | --- | --- |
|  | + | - | Total | + | - | Total | + | - | Total |
| CL cases | 32 | 22 | 54 | 29 | 25 | 54 | 23 | 31 | 54 |
| Non-CL cases | 1 | 49 | 50 | 1 | 49 | 50 | 2 | 48 | 50 |
| Total | 33 | 71 | 104 | 30 | 74 | 104 | 25 | 79 | 104 |
| Kappa index | 0,56<br>(0.42- 0.71)* |  |  | 0,51<br>(0.36-0.65) |  |  | 0.38<br>(0.23-0.52) |  |  |

Legend: DE: Direct Examination; HE: Histopathological Examination
CL Cases and non-CL cases were defined by PCR assay results
\*Confidence interval

### Serial Laboratory Testing

The diagnosis of CL can be challenging, and in some cases, additional laboratory tests on skin biopsy specimens are required to confirm the diagnosis. To improve diagnostic performance, a sequential (combined) analysis of tests was performed (Table 3).

**Table 3.** Diagnostic performance parameters for serial laboratory testing of DE plus anti-LACK IHC and HE plus anti-LACK IHC.

| Diagnostic tests | Sensitivity<br>(%) [CI95%]<br>n | Specificity<br>(%) [CI95%]<br>n | Accuracy<br>(%) [CI95%]<br>n |
| --- | --- | --- | --- |
| DE plus anti-LACK IHC | 72.2 [51 – 99]<br>(39/54) | 96 [70.8 – 100]<br>(48/50) | 83.7 [67-100]<br>(87/104) |
| HE plus anti-LACK IHC | 64.8 [45.2- 90.1]<br>(35/54) | 96 [70.8 – 100]<br>(48/50) | 79.8 [63.6-98.9]<br>(83/104) |

Legend: DE: Direct Examination HE: Histopathological Examination

The combination of DE with the anti-LACK IHC assay increased sensitivity from 53.7% to 72.2% (p = 0.23), slightly reduced specificity from 98% to 96% (p = 0.92), and improved overall accuracy from 75.0% to 83.7% (p = 0.48). Similarly, combining HE with anti-LACK IHC increased sensitivity from 42.6% to 64.8% (p = 0.12), maintained specificity at 96%, and improved accuracy from 68.3% to 79.8% (p = 0.33).

The results of the combined diagnostic methods (DE plus anti-LACK IHC and HE plus anti-LACK IHC) were cross-tabulated and are presented in Table 4. Comparison of the results of DE plus anti-LACK IHC with those of DE alone in relation to the CL case classification resulted in a Kappa coefficient of 0.68, indicating substantial agreement. Similarly, the cross-tabulation of HE plus anti-LACK IHC results yielded a Kappa coefficient of 0.60, corresponding to moderate agreement.

**Table 4.** Agreement analysis of the results of the diagnostic techniques used in this study.

|  | DE plus anti-LACK IHC |  |  | HE plus anti-LACK IHC |  |  |
| --- | --- | --- | --- | --- | --- | --- |
|  | + | - | Total | + | - | Total |
| CL cases | 39 | 15 | 54 | 35 | 19 | 54 |
| non-CL cases | 2 | 48 | 50 | 2 | 48 | 50 |
| Total | 40 | 5 | 104 | 37 | 67 | 104 |
| Kappa index | 0,68<br>(0.54- 0.81) |  |  | 0.60<br>(0.46-0.75) |  |  |

Legend: DE: Direct Examination HE: Histopathological Examination CL Cases and non-CL cases were defined by PCR assay results

## DISCUSSION

The diagnosis of CL remains challenging for clinicians, as its nonspecific clinical presentation may resemble a wide range of dermatological conditions, including allergic reactions to insect bites, stasis ulcers, sporotrichosis, leprosy, skin cancer, and lupus erythematosus [30]. When parasite burden is low, which is common in immunocompetent patients and chronic lesions, parasitological methods such as direct examination or culture often fail to detect the parasite.

In recent years, PCR assays targeting various *Leishmania* DNA sequences have demonstrated high sensitivity (75.7%–100%) and specificity (100%) [31]. In Brazil, however, PCR is not commercially available and is restricted to reference laboratories and research institutions, limiting patient access to accurate diagnosis. Conversely, IHC is routinely performed in public and private laboratories for the diagnosis of neoplastic, infectious, and parasitic diseases. In this context, IHC using mAbs against *Leishmania* antigens represents a promising approach to improve CL diagnosis and expand access to early treatment. In a previous study, an IHC assay based on a mAb against mitochondrial tryparedoxin peroxidase (mTXNPx) achieved a sensitivity of 85.7% in patients with CL and 37 controls with other diseases [19]. The present study developed and validated an IHC assay employing an anti-LACK mAb for CL diagnosis.

For several decades, the LACK protein has been investigated as a candidate for vaccine development, diagnostic testing, and as a therapeutic target [32–34]. Although the LACK sequences in *L. amazonensis*, *L. braziliensis*, and *L. guyanensis* are not completely identical, the anti-LACK mAb successfully recognized the native LACK antigen in soluble extracts and detected amastigotes in histological sections from hamsters experimentally infected with these main *Leishmania* species responsible for CL in Brazil. This finding is particularly important, as it indicates the potential applicability of this mAb across different endemic regions of the country.

Building on these findings, the IHC technique was validated using human skin lesion samples from patients with suspected CL. The anti-LACK IHC correctly identified 32 of 54 confirmed CL cases, resulting in a sensitivity of 59.3%, higher than that of direct examination (DE, 53.7%) and histopathology (HE, 42.6%), though the differences were not statistically significant (p = 0.70 and p = 0.22). Notably, disease duration did not significantly affect the sensitivity of anti-LACK IHC: sensitivity for participants with ≤3 months of illness was 50%, compared to 46.9% for those with 4–6 months of illness (p = 0.91, data not showed).

The sensitivity observed in this study for anti-LACK IHC (59.3%) was comparable to that reported for IHC assays using polyclonal sera and biotin-based detection systems, which ranged from 58.5% to 80% [13–17,35]. Similarly, Kenner et al. (1999) reported a sensitivity of 51% using an anti-*L. gerbilli* (G2D10) mAb with an avidin-biotin complex (ABC) detection system [36], whereas Lopes-Trujillo et al. (2021) achieved 94% sensitivity with a CD1a-specific mAb for 33 PCR-positive CL samples [37]. Shirian et al. (2014) also reported a sensitivity of 90% using mAbs against *L. major* and *L. tropica* in Iran [18]. Despite methodological heterogeneity among these studies, particularly regarding reference standards, sample sources, and IHC detection systems, the accumulated evidence supports IHC as a promising and sensitive approach for the diagnosis of CL.

The specificity of anti-LACK IHC (98%) was comparable to that of DE (98%) and HE (96%) (p = 0.92). Both anti-LACK IHC and DE yielded a single false-positive result, whereas HE produced two. The high specificity observed may be partially attributed to the use of Trilogy® and Bond Polymer Refine Red reagents, which minimized background staining and nonspecific labeling. Similarly, other studies have reported high specificity rates for IHC in CL diagnosis, ranging from 94.6% to 100% when using mAbs [17,19].

The agreement between anti-LACK IHC and the reference test (PCR–kDNA) was moderate (Kappa = 0.56). The lower positivity observed for anti-LACK IHC likely reflects the limited sensitivity of the technique, possibly due to the scarcity of amastigotes in the histological sections analyzed. It is well established that parasite burden decreases as lesion duration increases, particularly beyond six months of evolution [35–37]. Additionally, the tissue volume analyzed in IHC, DE, and HE is much smaller than that used for DNA extraction in PCR assays. In IHC and HE, only two sections of approximately 4 × 0.004 mm are examined, whereas DE uses eight imprints and PCR uses DNA extracted from tissue fragments of about 4 × 4 mm. Consequently, detecting *Leishmania* DNA by PCR is easier than visualizing amastigotes in histological sections or imprints. Supporting this, Sousa et al. (2014) reported positivity rates of 85.3% and 44% for a modified imprint method and conventional HE, respectively, for CL diagnosis [38].

The results of the present study demonstrated that combining DE with anti-LACK IHC increased sensitivity and accuracy without significantly reducing specificity (maintained at 96%). This sequential approach improved diagnostic accuracy from 75% to 83.7% (p = 0.48), reclassifying seven previously negative samples as CL-positive. These findings are consistent with Freire et al. (2022) [19], who reported an increase in CL positivity from 77.6% to 95.9% when combining DE with IHC. However, DE is not always available in pathology laboratories.

Similarly, combining HE with anti-LACK IHC reclassified 11 additional samples as CL-positive, improving diagnostic accuracy from 68.3% to 79.8% (p = 0.33) while maintaining specificity at 96%. Other authors have also observed improved positivity rates when HE and IHC were used together [14,17]. Since both methods employ histological sections, HE relying on direct staining and IHC on immunostaining, their combined use is feasible in pathology laboratories and may enhance diagnostic yield for CL.

In conclusion, the results of the present study do not support the use of anti-LACK IHC as a stand-alone diagnostic tool for CL. However, its application as a complementary test, particularly in combination with DE or HE, improves diagnostic performance. The development of additional mAbs targeting other *Leishmania* antigens should be encouraged to further optimize IHC-based diagnosis and support timely and accurate identification of CL in routine pathology practice.

## ACKNOWLEDGMENTS

We are grateful to Fernanda Oliveira Rodrigues for assistance with the histological sections. EO is grateful to CNPq-Brazil (Conselho Nacional de Desenvolvimento Científico e Tecnológico) for fellowships (Proc. 301555/2022-2)

## Supporting information captations

**Fig S1 captation:**

**Title:** Amino acid sequence of *Leishmania*-activated C kinase (A4HGX7_LEIBR)

**Legend:** The highlighted amino acids were used to design the final sequence of *Leishmania*-activated C kinase

**Tab S1 captation:**

**Title:** Optimization of immunohistochemistry protocol for detection of *Leishmania* using anti-LACK monoclonal antibody (anti-LACK IHC)

**Legend:** ¹Bond Polymer Refine Red Detection (Leica Microsystems, Newcastle, ENG)

